# *Anopheles gambiae* TEP1 forms a complex with the coiled-coil domain of LRIM1/APL1C following a conformational change in the thioester domain

**DOI:** 10.1101/550475

**Authors:** Marni Williams, Alicia Contet, Elena A. Levashina, Richard H. G. Baxter

## Abstract

The complement-like protein thioester-containing protein 1 (TEP1) is a key factor in the immune response of the malaria vector *Anopheles gambiae* to pathogens. Multiple allelic variants of TEP1 have been identified in laboratory strains and in the field, and are correlated with distinct immunophenotypes. TEP1 is tightly regulated by conformational changes induced by cleavage in a protease-sensitive region. Cleaved TEP1 forms a soluble complex with a heterodimer of two leucine-rich repeat proteins, LRIM1 and APL1C, and precipitates in the absence of this complex. The molecular structure and oligomeric state of the TEP1/LRIM1/APL1C complex is unclear. We have analyzed the stability of the cleaved form of four TEP1 alleles. Soluble TEP1 forms exhibit significant variation in stability from hours to days at room temperature. Stability is correlated with allelic variation within two specific loops in direct proximity to the thioester bond. The variable loops are part of an interface between the TED and MG8 domains TEP1 that protect the thioester from hydrolysis. Engineering specific disulfide bonds to prevent separation of the TED-MG8 interface stabilizes the cleaved form of TEP1 for months at room temperature. The C-terminal coiled-coil domain of the LRIM1/APL1C complex is sufficient to stabilize the cleaved form of TEP1 in solution but cleaved forms of disulfide-stabilized TEP1 do not interact with LRIM1/APL1C. This implies that formation of the TEP1_cut_/LRIM1/APL1C complex is dependent on the same conformational change that induces the precipitation of cleaved TEP1.

**Author Summary:** The mosquito *Anopheles gambiae* is the principal vector for malaria in Sub-Saharan Africa. A mosquito’s own immune system affects how readily it transmits disease. A protein in *A. gambiae* called TEP1 is responsible for targeting malaria parasites that traverse the mosquito’s midgut. TEP1 has multiple alleles and some are associated with a stronger immune response to malaria than others. How genetic variability in *TEP1* is linked to phenotypic diversity is not understood. We show that the variation between TEP1 alleles affects the stability of the protein in solution. We also show that the different TEP1 alleles have a wide range in stability of the protein, from hours to days. Engineering disulfide bonds into TEP1 can increase this stability to months. TEP1 activity *in vivo* is maintained by a complex of two leucine-rich proteins called LRIM1 and APL1C, which binds TEP1 through its C-terminal coiled-coil domain. We found that LRIM1/APL1C does not bind disulfide-stabilized TEP1, suggesting that LRIM1/APL1C binds to activated TEP1. This research advances our molecular understanding of a key immune response that affects the capacity of *A. gambiae* mosquitoes to transmit malaria.

## 1. Introduction

The mosquito *Anopheles gambiae* is the principal malaria vector in Sub-Saharan Africa. The immune response of *A. gambiae* is a significant factor influencing the vectoral capacity of mosquitoes to malaria parasites (genus *Plasmodium*). *Plasmodium* ookinetes invade and traverse the midgut epithelium, whereupon they face a robust complement-like immune response [1]. The complement-like factor thioester-containing protein 1 (TEP1) binds to the surface of *Plasmodium* ookinetes. TEP1-labeled ookinetes are targeted for killing by lysis and in some strains subsequent melanization.

TEP1 is a 160 kDa secreted glycoprotein comprised of eight fibronectin-fold domains called macroglobulin (MG) domains. In between MG7 and MG8 are two nested insertions of an eight stranded α-barrel domain (CUB) and the α-helical thioester domain (TED). The thioester domain contains a four amino acid motif – CGEQ – that forms a β-cysteinyl-γ-glutamyl thioester bond. The reactive thioester bond, a feature of the TEP protein family including α_2_-macroglobulins and complement factors, is protected from hydrolysis in the full-length protein by an interface formed between the TED and the MG8 domain [2]. A triangular arrangement of the TED, CUB and MG8 domains forms a ‘superdomain’ that is conserved between TEP1 and mammalian complement factor C3 [3, 4].

Activation of TEP1 requires cleavage in a protease-sensitive region located on an extended loop within the MG6 domain. The first six MG domains forma supermolecular ring, which is spanned by the protease-sensitive region, which connects to the MG7 domain and the TED-CUB-MG8 superdomain. Cleavage within the protease-sensitive region results in dissociation of the TED-MG8 interface, exposing the thioester bond for covalent reaction with substrates on pathogen surfaces, thereby labeling the pathogen with TEP1.

TEP1 is secreted into the mosquito hemolymph as a full-length protein; both full-length and cleaved TEP1 is detected in the hemolymph [5]. The cleaved form of TEP1 (TEP1_cut_) requires two leucine-rich repeat proteins, LRIM1 and APL1C, for stability in the hemolymph [6, 7]. In the absence of LRIM1 or APL1C, TEP1_cut_ precipitates over time both *in vitro* and *in vivo* [6, 8]. LRIM1 and APL1C consist of an N-terminal leucine-rich repeat (LRR) domain capped by a cysteine-rich motif. The two LRR proteins form a heterodimer via a C-terminal coiled-coil domain that contains a helix-loop-helix motif [9]. The LRIM1/APL1C heterodimer binds to TEP1_cut_ and stabilizes it in solution. TEP1_cut_ that is in complex with LRIM1/APL1C retains an intact thioester bond that allows for covalent attachment to a pathogen surface [8]. In the absence of LRIM1, the thioester bond hydrolyzes over time, resulting in non-functional precipitation of TEP1 [6, 8].

*A. gambiae* forms a species complex, comprising multiple morphologically identical forms across Sub-Saharan Africa [10]. *TEP1* is among the most polymorphic genes within *A. gambiae* genomes, with multiple distinct alleles that are correlated with susceptibility to *Plasmodium* infection [11]. TEP1*R1 is an allele associated with *A. gambiae* L3-5 laboratory strain, predominately found in the West African “M” molecular form of *A. gambiae*, now named *A. coluzzii* [12]. TEP1*R1 homozygous mosquitoes are refractory to the rodent malaria parasite *P. berghei* and have decreased susceptibility to the human malaria parasite *P. falciparum* [5, 13, 14]. *TEP1*S1* is an allele associated with *A. gambiae* G3 strain, found predominantly in the “S” molecular form of *A. gambiae*, which are susceptible to both *P. berghei* and *P. falciparum* infection. Other susceptible alleles named *TEP1*R2* and *TEP1*S2* are found in the laboratory 4arr strain and field isolates [11].

*TEP1* refractory and susceptible alleles are >90% identical with most variation being associated with variable loops within the TED, including two loops in direct proximity to the thioester bond. We previously reported that TEP1*S1_cut_ has a half-life of thioester hydrolysis *t*_1/2_ = 8.5 h, while TEP1*R1_cut_ has a half-life of thioester hydrolysis *t*_1/2_ = 6.5 days [8]. Replacing the TED of TEP1*R1 with that of TEP1*S1 reduced the half-life of thioester hydrolysis to almost the same as full-length TEP1*S_cut_, *t*_1/2_ = 12 h. This suggests that variation between TEP1 alleles within the thioester domain, specifically two hypervariable loops known as the pre-α4 and catalytic loop, respectively, contribute significantly to the stability of TEP1_cut_. Both alleles bind LRIM1/APL1C on a timescale consistent with their half-life for thioester hydrolysis.

Here we compared additional alleles and by mutation analysis identify specific residues within the loops of the TED that impact TEP1 stability. We further demonstrate that the coiled-coil domain of LRIM1/APL1C is sufficient to stabilize TEP1_cut_ in solution. Finally, we show that engineered disulfides within the TED-CUB-MG8 superdomain can stabilize TEP1_cut_ similar to LRIM1/APL1C. These results suggest that the formation of the TEP1_cut_/LRIM1/APL1C complex requires a conformational change involving dissociation of the TED-MG8 interface, the same conformational change that leads to thioester hydrolysis and precipitation of TEP1cut in the absence of LRIM1/APL1C.

## Results

### Rate of thioester hydrolysis for TEP1 alleles

Polymorphisms between TEP1*R1 and TEP1*S1 are concentrated within and adjacent to the TED, and specifically in three hypervariable loops within the TEP. Two of these loops, the catalytic loop and the pre-α4 loop, are in close proximity to the thioester bond and form part of the TED-MG8 interface. The third loop, the b-hairpin loop, lies on the opposite side of the TED, surface-accessible and near the TED-MG8 interface. We previously showed that a chimeric protein replacing the TED of TEP1*R1 with TEP1*S1 had the same stability as TEP1*S1, but this does not determine which of the three loops are responsible for this variable stability.

Blandin et al. (2009) reported two additional alleles, *TEP1*S2* and *TEP1*R2*, that are virtually identical to *TEP1*S1* and *TEP1*R1*, respectively, within the TED (Fig. 1). The most notable difference between TEP1*S1 and TEP1*S2 is that the TEP1*S2 β-hairpin motif is identical to TEP1*R1 and TEP1*R2. Hence we used TEP1*S2 to test whether the R1/R2 β-hairpin motif has a stabilizing effect on TEP1_cut_.

**Figure 1.**
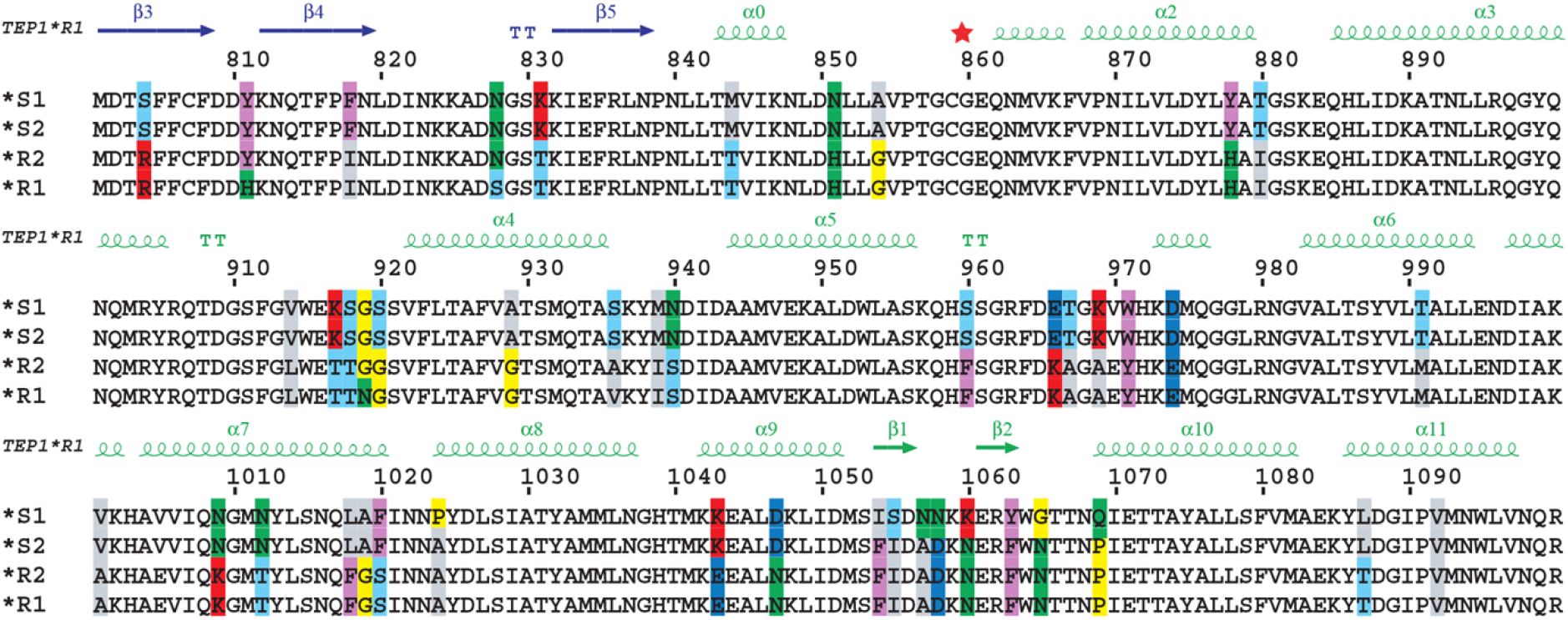
Multiple sequence alignment of TEP1 forms TEP1*S1, TEP1*S2, TEP1*R2 and TEP1*R1 residues 800-1100, including the hypervariable TED pre-α4 loop (917-920), catalytic loop (966-971) and β-hairpin (1054-1069). Adapted from Blandin et al. (2009) [11].

We produced TEP1*S2, performed limited proteolysis and measured the half-life of thioester hydrolysis in three independent experiments (Table 1). We observed that TEP1*S2_cut_ had a half-life of thioester hydrolysis *t*_1/2_ = 4.2 ± 0.2 h, half that of TEP1*S1_cut_ *t*_1/2_ = 8.5 ± 0.2 h. Hence, the β-hairpin motif is not responsible for the stabilization of TEP1*R1 relative to TEP1*S1, consistent with its location opposite the thioester bond and the TED-MG8 interface [8, 15].

**Table 1:** Rate of thioester hydrolysis of TEP1_cut_

| TEP1 form | $t_{1/2}$ (h) |
| --- | --- |
| *S2 | $4.2 \pm 0.2$ |
| *S1 | $8.5 \pm 0.2$ |
| *R2 | $9.6 \pm 0.4$ |
| **\*R1** | **$156 \pm 12$** |
| *R1 S1<sub>cat</sub> | $9 \pm 2$ |
| **\*R1 N919G** | **$55 \pm 10$** |

TEP1*R2 is a chimeric form whose N-terminal fragment is similar to S1/S2, while the C-terminal fragment is almost identical to TEP1*R1 except for a single residue in the TED pre-α4 loop, TEP1*R1 Asn 919, which is glycine in TEP1*R2, TEP1*S1 and TEP1*S2. To test whether the R1 mutation G919N has a stabilizing effect on TEP1_cut_, we produced TEP1*R2, performed limited proteolysis and measured the half-life of thioester hydrolysis. We observed that TEP1*R2_cut_ had a half-life of thioester hydrolysis *t*_1/2_ = 9.6 ± 0.4 h, comparable to that of TEP1*S1_cut_, and 16 times less than that of TEP1*R1_cut_ *t*_1/2_ = 156 ± 12 h. This suggests that the differences between TEP1*R1 and TEP1*R2, specifically the point mutation N919G, have a significant effect on TEP1_cut_ stability.

To confirm the role of Asn 919, we made the point mutation N919G in TEP1*R1 and measured the stability of TEP1*R1-N919G_cut_. This single mutation reduced the half-life of thioester hydrolysis three times, TEP1*R1-N919G_cut_ *t*_1/2_ = 55 ± 10 h. The differences between TEP1*R1-N919G_cut_ and TEP1*R2 that contribute to further destabilization of TEP1*R2_cut_ lie outside of the TED and MG8 domains. The closest TEP1*R2 mutations relative to TEP1*R1 (i.e. conserved with TEP1*S1, TEP1*S2) are H811Y and S828N in the CUB domain.

The catalytic loop is the third hypervariable loop in the TEP1 TED, and is in direct proximity to the thioester bond. That catalytic loop is hypervariable between TEP1*S1/TEP1*S2 and TEP1*R1/TEP1*R2, hence we expected it to directly impact the stability of TEP1_cut_. We therefore made the mutation TEP1*R1-S1_cat_, substituting residues 966–971 from KAGAEY in TEP1*R1 to ETGKVW. The cleaved form of TEP1*R1-S1_cat_ has a half-life of thioester hydrolysis *t*_1/2_ = 9.1 ± 1.9 h, comparable to that of full-length TEP1*S1. Hence the catalytic loop alone is sufficient to explain the destabilization of TEP1*S1_cut_ relative to TEP1*R1. Put another way, the unusual stability of TEP1*R1 compared to TEP1*S1, TEP1*S2 and TEP1*R2 is the result of specific mutations in both the pre-α4 and the catalytic loop with some contribution of TEP1*R1-specific mutations outside the TED-MG8 domains.

### Stabilization of TEP1_cut_ by engineered disulfide bonds

Following cleavage within the protease-sensitive region, TEP1_cut_ slowly converts from a state that contains an intact thioester to one in which the thioester is hydrolyzed and which precipitates from solution. Cleavage of the protease-sensitive region occurs in a separate location from the TED-MG8 interface, the MG6 domain. Yet the TED-MG8 interface must separate for the thioester bond to bind to a substrate. Association of LRIM1/APL1C with TEP1 forestalls hydrolysis of the thioester, but how LRR complex binding stabilizes TEP1_cut_ is unknown. We tested whether LRIM1/APL1C binding is separable from thioester hydrolysis and concomitant precipitation of TEP1_cut_ by stabilizing the TED-MG8 interface with engineered disulfide bonds.

Guided by the crystal structure of TEP1*S1, we identified residues at the TED-MG8 and TED-CUB interfaces where mutation to cysteine would generate a disulfide bond with minimal perturbation of the structure [8]. We generated three disulfide mutants of TEP1*S1, varying the location of the disulfide between the TED and CUB/MG8 domains (Fig. 2A). First, we joined TED residue K973 within the catalytic loop to residue Q1228 in the MG8 domain (K973C/Q1228C). This had the effect of increasing the half-life for thioester hydrolysis to *t*_1/2_ = 4.0 days (Fig. 2B). Second, we joined TED residue V1102 to residue M801 in the CUB domain (M801C/V1102C). This change increased the half-life for thioester hydrolysis to *t*_1/2_ = 44 days. Third, we joined TED residue K969 within the catalytic loop to a residue in the MG8 domain Y1275 (K969C/Y1275C). This resulted in a half-life for thioester hydrolysis of *t*_1/2_ > 100 days. This confirms that precipitation of TEP1_cut_ occurs after separation of the TED-MG8 domain, and preventing this conformational change can effectively stabilize TEP1_cut_ indefinitely.

**Figure 2:**
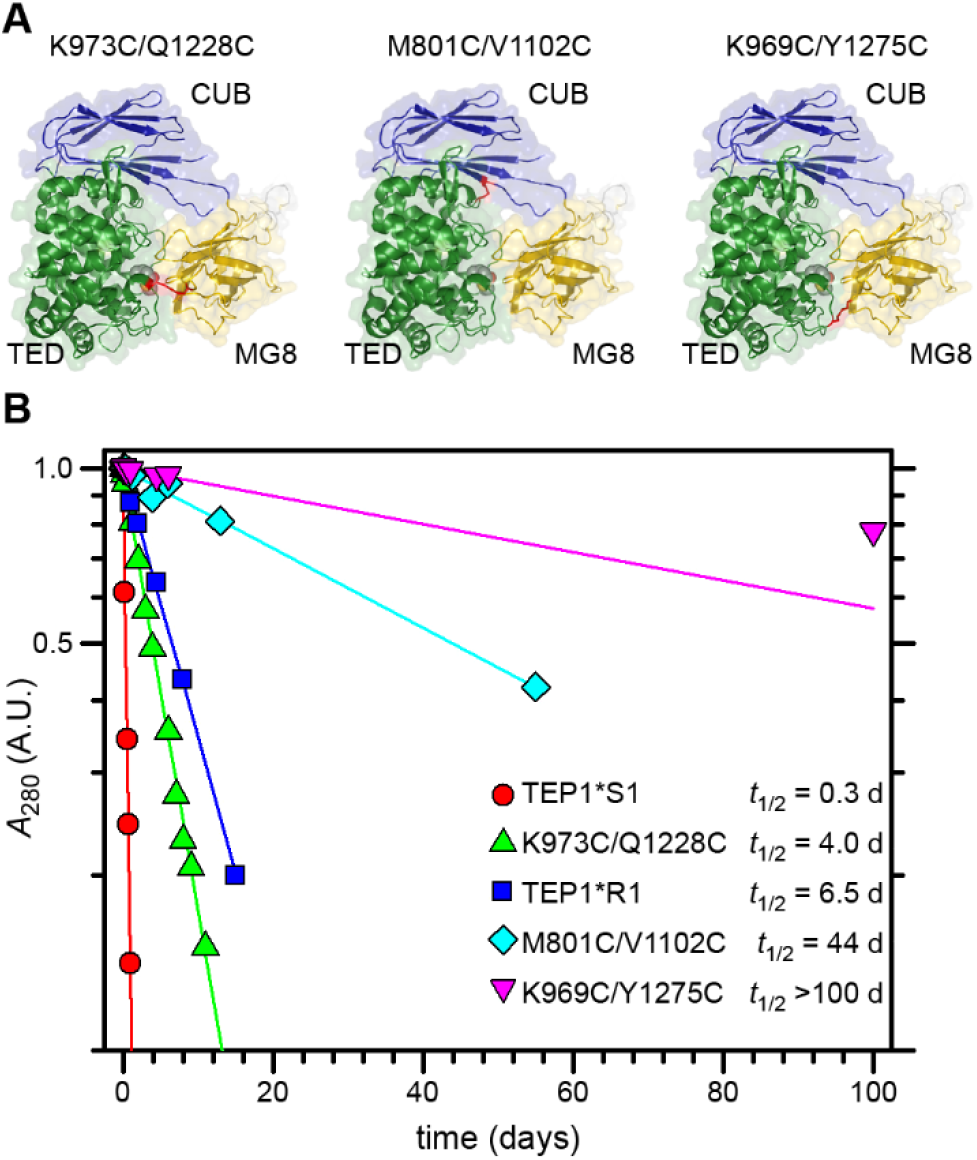
(A) Model of TED (green), CUB (blue) and MG8 (yellow) domains for TEP1*S1, with engineered disulfide bonds (red) K973C/Q1228C, M801C/V1102C and K969C/Y1275C. (B) Rate of thioester hydroylsis for cleaved forms of TEP1*S1, TEP1*R1, and three engineered disulfide mutants of *TEP1*S1*.

### Stabilization of TEP1_cut_ by LRIM1/APL1C coiled-coil domain

The LRIM1/APL1C heterodimer has three discrete, pseudosymmetric structural elements (Fig. 3A). Each protein has an LRR domain capped by an LRRCT helix, followed by a short coiled-coil domain, an intermolecular disulfide, then a helix for LRIM1 and a flexible region for APL1C. Both proteins end with a C-terminal coiled-coil domain that contains a helix-loop-helix (HLH) motif. Multiple groups have shown that full-length LRIM1/APL1C stabilizes TEP1_cut_ *in vitro* and *in vivo* [6–9, 16]. However, the LRIM1 and APL1C LRR domains were previously shown to be insufficent to stabilize TEP1*R1_cut_ after treatment with methylamine (MeNH_2_) to chemically inactivate the thioester bond, or to stabilize endogenous TEP1_cut_ *in vivo* [9]. Povelones et al. (2011) expressed LRIM1/APL1C with internal deletions of the coiled-coil domain [16]. These constructs were unable to bind cleaved TEP1. In this study however, TEP1_cut_ was produced by co-expression of separate N- and C-terminal fragments. As the structure of the TEP1 MG6 domain is formed by the intertwining of both N- and C-terminal fragments; co-expressing both fragments separately may result in misfolding of TEP1.

**Figure 3.**
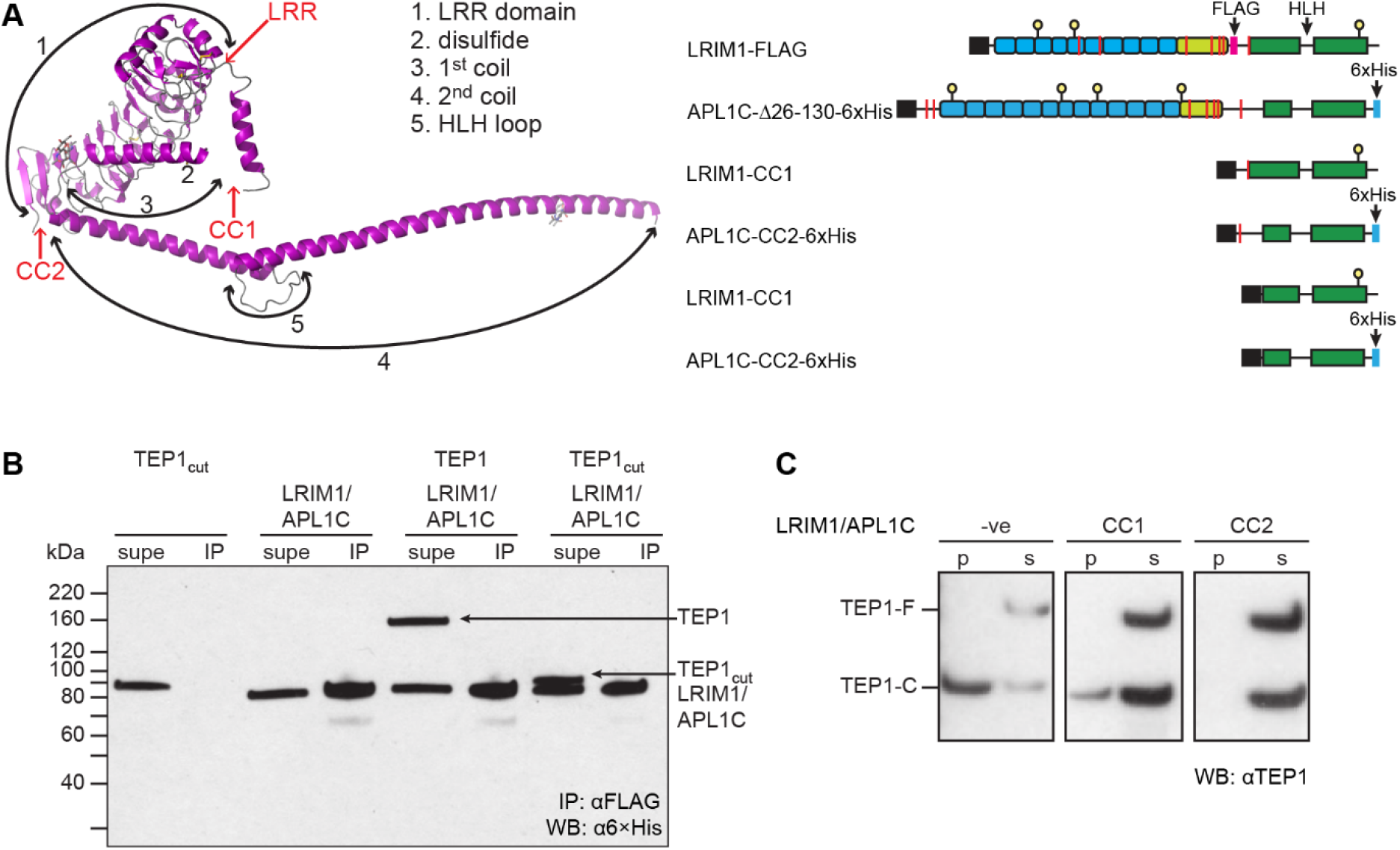
(A) Model of the domain structure of LRIM1/APL1C (LRIM1 only, magenta) indicating LRR domain and location of C-terminal truncations CC1 and CC2. A schematic diagram of LRIM1-FLAG/APL1C-Δ26-130-6xHis construct and coiled-coil domain constructs CC1 and CC2, Coiled-coil shown in green, cysteines as red bars, N-linked glycosylation site as yellow ball, 6xHis tag in blue. (B) a6×His Western blot of supernatant (supe) and αFLAG immunoprecipitate (IP) of TEP1*S1-K969C/Y1275C seven days post-cleavage. The first sample is wt TEP1*S1 immediately following cleavage. Both uncleaved and cleaved TEP1 remain in the supernatatant and do not co-immunoprecipitate with LRIM1/APL1C. (C) αTEP1 Western blot of precipitate (p) and soluble (s) fractions for TEP1*S1 full-length (TEP1-F) and cleaved form (TEP1-C) after 24 h incubation with LRIM1/APL1C-CC1 and LRIM1/APL1C-CC2.

We therefore sought to test two hypotheses, (i) that LRIM1/APL1C binding is associated with conformational change of the TED-MG8-CUB superdomain by exploiting our disulfide-stabilized mutants, and (ii) that the coiled-coil domain of LRIM1/APL1C is sufficient to stabilize TEP1_cut_. First, we examined the ability of engineered disulfide mutants of TEP1*S1 to bind the LRIM1-FLAG/APL1C-Δ26-130-6xHis construct previously found to stabilize TEP1*S1 [8, 9]. Then, to show that the C-terminal coiled-coil domain is the site of TEP1 binding we produced two C-terminal coiled-coil constructs of LRIM1/APL1C. LRIM1/APL1C-CC1 comprises all C-terminal residues from the intermolecular cysteines LRIM1 C352 and APL1C C551. LRIM1/APL1C-CC2 comprises the C-terminal coiled-coil domain including the helix-loop-helix motif.

Despite the dramatically increased stability of the disulfide-engineered TEP1*S1 mutants, none were stabilized by or co-immunoprecipitated with LRIM1-FLAG/APL1C-Δ26-130-6xHis (Fig. 3B). Wild-type TEP1*S1_cut_ does not IP with αFLAG immediately following cleavage. Neither full-length or cleaved TEP1*S1-K969C/Y1275C co-immunoprecipitated with LRIM1-FLAG/APL1C-Δ26-130-6xHis seven days post-cleavage. Three months (100 days) later, the result of TEP1*S1-K969C/Y1275C αFLAG co-IP with LRIM1-FLAG/APL1C-Δ26-130-6xHis was unchanged (data not shown).

We then tested whether the coiled-coil domain of LRIM1/APL1C was sufficient to prevent the precipitation of TEP1_cut_. When incubated with a mixture of TEP1*S1 and TEP1*S1_cut_, both LRIM1/APL1C-CC1 and LRIM1/APL1C-CC2 were able to stabilize TEP1*S1_cut_ in solution (Fig. 3C). This demonstrates that the C-terminal coiled-coil domain of LRIM1/APL1C is necessary and sufficient to stabilize TEP1_cut_.

## Discussion

TEP1 is a key antiparasitic factor in *A. gambiae*, and allelic variation in TEP1 significantly influences susceptibility of *A. gambiae* to *Plasmodium*. Most of the variation between alleles is within the thioester domain, specifically three loops of which two – the pre-α4 loop and the catalytic loop – are proximal to the thioester bond and form part of the TED-MG8 domain interface. Variation in the pre-α4 and the catalytic loop can potentially affect the reactivity and substrate specificity of the thioester as well as the overall stability of the protein in solution. However, these inter-related properties complicate understanding the molecular mechanism of TEP1.

We previously reported a significant difference in stability for the cleaved alleles of TEP1*S1 and TEP1*R1 [8]. Here we report that genetic polymorphism in the TED domain of *TEP1* shapes its function by regulating protein stability. Interestingly, TEP1*R1, the only allele that confers mosquito resistance to *Plasmodium* infection, has an exceptionally long lifetime. Other forms display more subtle variation, ranging from 9.2 h for *R2 to 4.2 h for *S2. Whether the increased stability of *R1 directly causes its antiparasitic activity remains to be demonstrated.

The increased stability of TEP1*R1 requires *both* changes in the catalytic loop sequence 966–971 from ETGKVW (S) to KAGAEY (R) and in the pre-α4 loop sequence 917-920 from KSGS (S) to TTNG (R), especially the mutation G919N. This is drawn from the fact that reversion of either 966-971 of 919 from R to S significantly decreases stability of cleaved TEP1. These changes are synergistic, because TEP1*R2 which is identical with TEP1*R1 in both these loops but for G919N is only marginally more stable than TEP1*S1. Yet interactions outside the TED-MG8 domain contribute to stability, because the mutant TEP1*R1-N919G is identical to TEP1*R2 in both the TED and MG8 domains yet is still more stable (*t*_1/2_ = 2.3+0.4 days vs. 9.6+0.4 h).

An open question is whether the conformational change in TEP1_cut_ that permits LRIM1/APL1C binding is separable from that which induces thioester hydrolysis and precipitation. We sought to address this by engineering specific disulfide bonds between domains. We were able to stabilize TEP1*S1_cut_ for months with engineered disulfides from the TED to either the MG8 or CUB, but in neither case did the cleaved protein interact with LRIM1/APL1C. Hence, it appears the same conformation involving separation of the TED from CUB and MG8 is responsible for both LRIM1/APL1C binding and TEP1 aggregation and precipitation.

Figure 4 illustrates the location of the three engineered disulfides to the TED-MG8 interface and the thioester bond. The first disulfide replaces K793 directly adjacent to the catalytic histidine of the TED catalytic loop and Q1228 at the top of the MG8 loop in the center of the interface near M1231, which directly abuts the thioester. Interestingly, this was the least stabilizing disulfide, less stable than TEP1*R1, suggesting that securing the catalytic loop to the top of the TED-MG8 interface does not prevent the interface from separating beneath it, possibly due to perturbation of the domains MG2 and MG6 that lie below. In comparison, the disulfide replacing K969 in the catalytic loop andY1275, one of the aromatic residues that ring the thioester bond, is indefinitely stabilizing.

**Figure 4.**
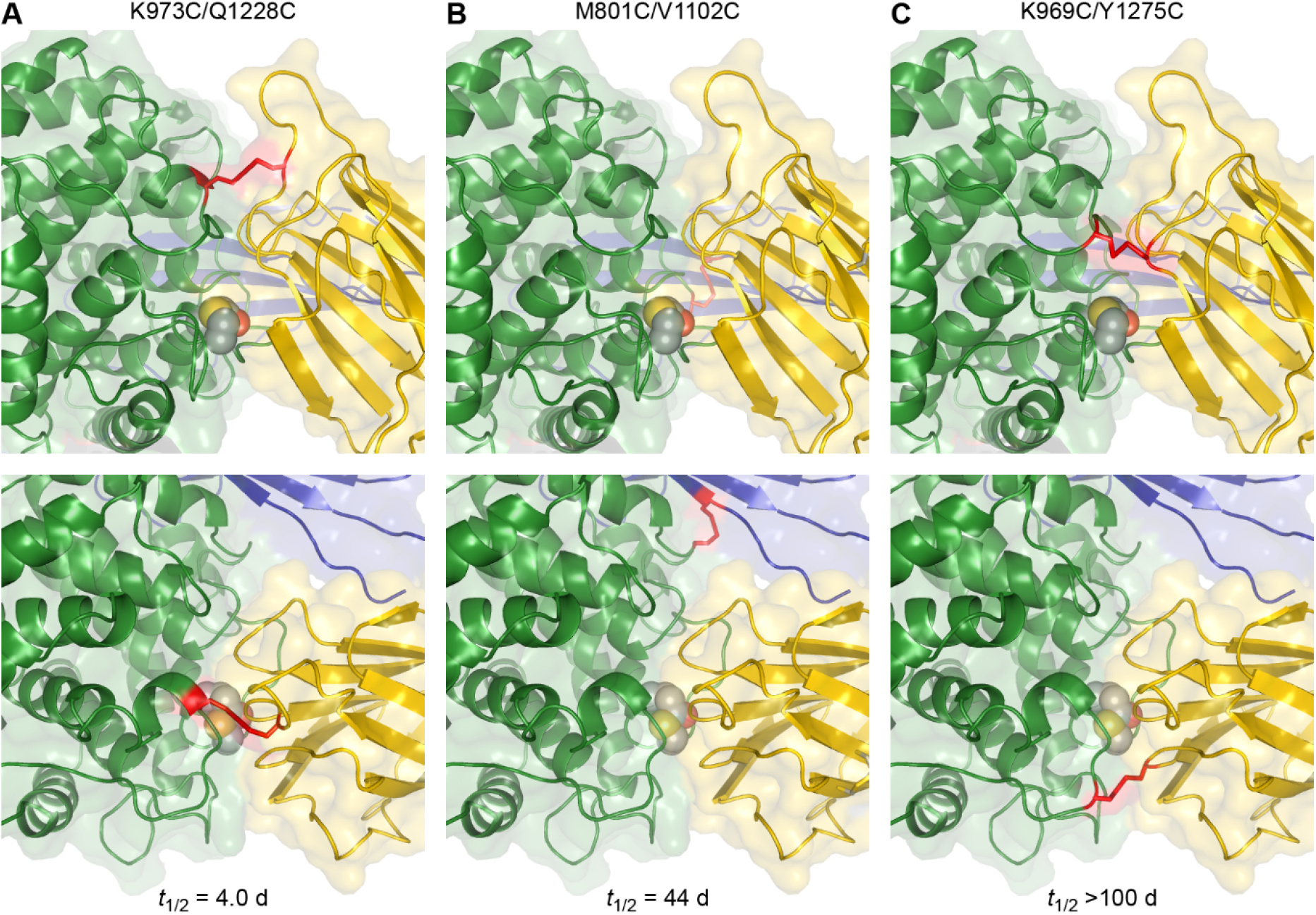
Front (upper) and top (lower) views of the TED-MG8 domain interface of TEP1*S1 with modeled engineered disulfides K973C/Q1228C, M801C/V1102C and K969C/Y1275C.

Surprisingly, the third disulfide engineered between TED V1102 and CUB M801 is a highly stabilizing mutant even though it does not involve the TED-MG8 interface. In the conversion of complement C3 to C3b and C5 to C5b, the CUB domain undergoes a large motion along with the TED, hence locking the relative position of these domains may equally serve to prevent separation of the TED-MG8 interface.

In vertebrate complement factors the anaphylatoxin domain serves as a molecular wedge between the MG3 and MG8 domain, and we had speculated that LRIM1/APL1C might form an analogous complex following cleavage of TEP1. The fact that TED-MG8 engineered disulfides prevent LRIM1/APL1C binding run counter to this hypothesis. It suggests that the LRIM1/APL1C coiled-coil domain has a cryptic binding site somewhere on the TED, CUB and/or MG8 that is only exposed by dissociation of this superdomain. Povelones et al. (2011) found that both N- and C-terminal fragments were required for TEP1 to interact with LRIM1/APL1C [16]. Considering the C-terminal coiled coil domain is 17 nm end-to-end, longer than TEP1 itself, it is entirely possible that additional interactions could involve the N-terminal fragment of TEP1.

The LRIM1/APL1C heterodimer has long been known to stabilize TEP1_cut_, and it has been shown that the LRR domains alone or as heterodimers with deletions in the coiled-coil domain are insufficient to stabilize TEP1_cut_ [9, 16]. Here, we specifically purified heterodimers of only the coiled-coil domains of LRIM1/APL1C and show they are sufficient to stabilize TEP1*S1_cut_, confirming that the coiled-coil domain is both necessary and sufficient to stabilize TEP1_cut_. This is significant because the coiled-coil domain is shared among the related genes APL1A^1^ and APL1B, which also form a complex with LRIM1 [17].

As presently stands, the inability to isolate a stable complex of TEP1_cut_ and LRIM1/APL1C in solution prevents structural studies that require a homogeneous and monodisperse sample. Additional engineering to solubilize TEP1_cut_ or prevent its aggregation, the identification of orthologs or paralogs for the TEP1_cut_/LRIM1/APL1C complex, or approaches for the analysis of heterogeneous specimens are necessary to define the molecular basis of LRIM1/APL1C regulation of TEP1 activation.

## Materials and Methods

### Molecular cloning

DNA for *TEP1*R1* and *TEP1*S1* was obtained from cDNA clones or by total gene synthesis as previously described [8, 15]. DNA for *TEP1*R2* and *TEP1*S2* were generated by total gene synthesis from sequences in the 4arr laboratory strain [11]. All *TEP1* sequences were subcloned into pFastbac1 with a C-terminal 6×His tag. Additional mutations within the thioester domain were introduced by QuickChange site-directed mutagenesis (Stratagene). *LRIM1* and *APL1C* were subcloned into the pFastbac-Dual vector with C-terminal 6×His tag on APL1C. Construction of LRIM1-FLAG/APL1C-D26-130-6xHis has been previously described [8, 9]. Truncations of the LRR and coiled-coil domains were introduced by site-directed mutagenesis.

### Protein expression and purification

All TEP1 and LRIM1/APL1C constructs were expressed using the baculovirus expression system using T.ni cells in ESF-921 media (Expression Systems LLC). TEP1 was expressed for ~60 h at 27°C, LRIM1/APL1C for ~48 h at 27°C. Conditioned medium was concentrated and diafiltrated with 250 mM NaCl, 20 mM Tris-HCl pH 7.8 using Centramate tangential flow filtration system (Pall Biosciences). The diafiltrated medium was loaded onto 5 ml Talon resin (Clontech), washed with 10 CV of 250 mM NaCl, 20 mM Tris-HCl pH 7.8 and eluted by a 20 ml gradient to 250 mM imidazole.

TEP1 eluate from Talon resin was immediately desalted on a HiTrap 26/10 column (GE Healthcare) equilibrated with 100 mM NaCl, 20 mM Tris-HCl pH 8.0, loaded onto a MonoQ column (GE Healthcare) and eluted with a linear gradient from 100–600 mM NaCl. Further purification was achieved by gel filtration (Superdex 200) equilibrated with 100 mM NaCl 20 mM Na-Hepes pH 7.5 and cation exchange on a MonoS column. LRIM1/APL1C eluate from Talon resin was desalted into 50 mM NaCl, 20 mM Tris-HCl pH 8.5 prior to MonoQ chromatography followed by final purification on gel filtration (Superdex 200) equilibrated with 100 mM NaCl 20 mM Na-Hepes pH 7.5.

### Limited Proteolysis of TEP1

Purified TEP1 was cleaved using bovine pancreatic trypsin (Sigma) at a 1:20 molar ratio to TEP1 in 0.2 M NaCl and 20 mM HEPES pH 7.5. Samples were incubated for 5 min at 37°C, then placed on ice and diluted 2-fold with 20 mM HEPES pH 7.5, 0.2 mM Leupeptin hemisulfate (Sigma), and 0.2 mM soybean trypsin-chymotrypsin inhibitor. Samples were immediately repurified on a Mono S 5/50 cation exchange column (GE Healthcare).

### Thioester hydrolysis precipitation assay

Following limited proteolysis and re-purification, TEP1 samples were concentrated to an OD_280_ of 0.5–1.0 and stored at 20°C. Samples and matching blank (filtrate from concentration) were centrifuged at 17,000×*g*, 20°C for 10 min and *A*_280_-*A*_330_ recorded. Separate time points are all derived from the same protein batch. Half-lives were calculated from samples with a decay to <25% initial value and results derived from three independent experiments.

### Co-immunoprecipitation and Western Blotting

Rabbit polyclonal antibodies for TEP1*R1 have been previously described [9]. 10 μg of protein mixtures was diluted to 1 ml in IP buffer (50 mM Tris-HCl pH 7.8, 100 mM NaCl, 2 mM EDTA, 0.1 μg/ml BSA, 0.1% Tween-20). IP was performed with αFLAG-M2 agarose (Sigma). Beads were washed twice each with 50 mM Tris-HCl pH 7.8 ± 0.5 M NaCl and eluted by incubation with 2X Laemmli buffer. SDS/PAGE was run with 4-20% minigels, transferred to nitrocellulose and Western blotting performed with monoclonal α6xHis/HRP (Clontech) or αTEP1.

## Acknowledgements

This work was supported in part by the National Institute of General Medical Sciences (1R01GM114358) of the National Institutes of Health to RHGB.

## Author Contributions

Conceptualization, M.W., A.C., E.A.L. and R.H.G.B.; Methodology, M.W., A.C., and R.H.G.B.; Investigation, M.W. and A.C.; Writing – Original Draft, E.A.L. and R.H.G.B.; Writing – Review and Editing, E.A.L. and R.H.G.B.; Funding Acquisition, E.A.L. and R.H.G.B.; Resources, E.A.L. and R.H.G.B.; Supervision, R.H.G.B.

